# Secondary nucleation drives polymorph diversity in hIAPP amyloids

**DOI:** 10.64898/2026.08.20.745575

**Authors:** Mikołaj I Kuska, Łucja Kozicka, Sayan Prodhan, Dylan Valli, Michał Maj

## Abstract

Amyloid fibrils are implicated in a myriad of human diseases. A striking observation is that fibrils extracted from diseased tissues are characterized by a restricted set of folds unique to the specific pathology. In contrast, fibrils grown *in vitro* exhibit extensive structural diversity, suggesting that specific environmental and biochemical mechanisms *in vivo* enforce structural selectivity. Here, we combine two-dimensional infrared (2D IR) spectroscopy and cryo-electron microscopy (cryo-EM) to investigate the mechanisms governing polymorph formation in the human Islet Amyloid Polypeptide (hIAPP). We demonstrate that 2D IR can resolve populations of distinct polymorphs identified by cryo-EM, enabling rapid label-free screening of conditions prior to labor-intensive microscopy screening. We find that conditions favoring secondary nucleation, such as high protein concentration, increase polymorphic diversity. Crucially, cryo-EM reveals that formed by secondary nucleation do not structurally replicate the parent template. Finally, by selectively inhibiting secondary nucleation using the C-terminal domain of the DNAJB6 chaperone, we steer aggregation toward a monomorphic state. These findings highlight the critical role of molecular chaperones in fibril polymorph selection.

## INTRODUCTION

Amyloid diseases arise when proteins or peptides access misfolded conformations that self-assemble into highly ordered, cross-*β* fibrils and deposit in tissues and organs.^1,2^ Amyloid deposits are implicated in numerous disorders, ranging from neurodegenerative diseases such as Alzheimer’s disease to metabolic disorders such as type 2 diabetes, where human islet amyloid polypeptide (hIAPP, amylin) is the principal component of islet amyloid.^3–5^

A central complexity in amyloid biology is that a single polypeptide sequence can adopt multiple distinct fibril structures known as polymorphs. These polymorphs can differ in protofilament fold, inter-protofilament interactions, and the presence of bound cofactors. Such structural differences are associated with altered stability, toxicity, seeding efficiency, and pathology. ^6–10^ Fibril elongation transmits structural information through templated addition at fibril ends, which provides a conceptual basis for prion-like inheritance of conformers.^9,11^

Over the last decade, cryo-electron microscopy (cryo-EM) has provided near-atomic structures for a growing set of disease-associated amyloids and highlighted differences between *ex vivo* (patient-derived) and *in vitro* fibrils.^12–14^ Despite *in vitro* grown tau fibrils yielding almost 80 different polymorphs,^15^ *ex vivo* filaments from the human brain are uniquely defined and disease-specific, such that Alzheimer’s disease tau filaments are structurally different from other neurodegenerative disorders, such as Pick’s disease, for instance.^16 12,17^ Such observation is also true for amyloid-*β* peptide, ^14,18^ and *α*-synuclein.^13^ Nevertheless, *in vitro* seeded amplification is not guaranteed to preserve these patient-derived structures: seeded assembly of recombinant *α*-synuclein with brain-derived Multiple System Atrophy (MSA) seeds produces filaments that differ from the seeds. This implies that factors beyond protein sequence and seed conformation influence structural propagation.^19^

These discrepancies pose a practical challenge because differences in polymorph composition can alter aggregation kinetics, toxicity, and inhibitor sensitivity, complicating comparisons between studies and limiting their disease relevance.

For hIAPP, multiple *in vitro* cryo-EM structures have revealed extensive polymorphism and provided a basis for inhibitor design.^20–23^ Moving closer to clinical relevance, Cao *et al*. extracted hIAPP fibrils from a donor with type 2 diabetes, used them to seed synthetic hIAPP, and solved four cryo-EM polymorphs whose similarity to unseeded *in vitro* structures ranged from comparable to negligible - highlighting that patient-derived material strongly influences the resulting fibril structures.^24^ Most recently, direct cryo-EM determination of pancreatic hIAPP fibrils extracted from donors with type 2 diabetes revealed a morphology with an Ω-shaped protofilament fold that differs from known *in vitro* and seeded structures and displays additional densities consistent with ligand binding. ^25^ Together, these studies indicate that while many folds are accessible *in vitro*, the *in vivo* environment imposes strong selection pressures that yield a restricted set of dominant structures. Reproducibly controlling hIAPP polymorph distributions would help identify the factors that drive this selection.

Understanding this structural disparity requires considering that amyloid formation is not a single pathway but a network of microscopic steps– primary nucleation, elongation, fragmentation, and secondary nucleation–whose relative weights can shift with solution conditions and surfaces. ^26–29^ Secondary nucleation, in which fibril surfaces catalyze the formation of new nuclei, is a particularly powerful autocatalytic amplifier because it can generate large numbers of new fibrils once a fibril surface area is present.^27,28^ For IAPP, surface-catalyzed secondary nucleation has been implicated in the formation of toxic aggregates.^30^ Unlike end-templated elongation, secondary nucleation is a *de novo* nucleation event occurring on a heterogeneous and evolving surface; therefore, it may bias nucleation without enforcing faithful replication of the parent fibril architecture. Consistent with this idea, time-dependent cryo-EM has shown that IAPP fibril populations can undergo structural evolution over the course of assembly, with new polymorphs appearing and others disappearing as reactions progress.^31^ This behavior was proposed to reflect changes in the relative contributions of primary nucleation, secondary nucleation, and elongation during the aggregation process - leaving open the key question of whether secondary nucleation propagates the parent fibril structure or generates structurally distinct polymorphs.

Answering this requires methods that connect kinetics to structure. Cryo-EM provides near-atomic models but remains low-throughput for mapping polymorphs across broad conditions. Two-dimensional infrared (2D IR) spectroscopy offers a complementary approach: it is rapid, sensitive to vibrational couplings within the Amide I manifold that report on *β*-sheet geometry, and can resolve heterogeneous mixtures through distinct spectral signatures. ^32–35^ An efficient strategy may therefore be to use 2D IR for ensemble screening and cryo-EM for structural validation.

Here, we connect the microscopic kinetics of hIAPP aggregation to the resulting fibril structures using label-free 2D IR spectroscopy and cryo-EM. We first show that 2D IR spectroscopy resolves the distinct vibrational signatures of coexisting polymorphs, providing a rapid measurement of structural distributions. By varying the initial peptide concentration, we alter the relative contribution of secondary nucleation pathways during assembly and observe a corresponding increase in polymorphic diversity. To explicitly test the role of this pathway, we introduce the C-terminal domain of the DNAJB6 chaperone (DNAJB6-CTD) to selectively suppress surface-catalyzed nucleation. ^36^ Blocking this process restricts the hIAPP population to a single dominant fibril architecture. Together, these results demonstrate that secondary nucleation is a primary driver of structural heterogeneity in IAPP and that chaperone proteins can physically dictate polymorph selection.

## METHODS

### Sample preparation

hIAPP was synthesized on a 0.1 mmol scale using standard 9-fluorenylmethyloxycarbonyl (Fmoc) solid-phase peptide synthesis on a Biotage Initiator+ Alstra peptide synthesizer. Tentagel R RAM resin was used to obtain the C-terminal amide. Pseudoproline dipeptide derivatives were incorporated to facilitate coupling of difficult regions according to established IAPP synthesis protocols. ^37^ The final product was cleaved from the resin using a cocktail of 95% TFA, 2.5% H_2_O, and 2.5% triisopropylsilane for 2 h. The crude peptide was precipitated with cold diethyl ether, redissolved in H_2_O, and lyophilized. The peptide was oxidized for 24 h to form the disulfide bridge between Cys2 and Cys7 by dissolving in 60% DMSO with 8% acetic acid aqueos solution. Purification was performed using reverse-phase HPLC (Isera C18 preparative column, 250 mm *×* 20 mm) with a gradient elution composed of buffer A (H_2_O with 0.045% HCl) and buffer B (80% acetonitrile with 0.045% HCl). After lyophilization, the peptide was resuspended in 50% hexafluoroisopropanol (HFIP) and 5% acetic acid and purified a second time. The peptide mass was confirmed using MALDI-FTICR-MS. The pure peptide was monomerized by resuspending in 100% HFIP and filtering through a 0.22 *µ*m PTFE filter. The concentration was determined from the absorbance at 280 nm using the extinction coefficient of tyrosine (*ε*_280_ = 1490 M^*−*1^ cm^*−*1^). The peptide was aliquoted and lyophilized to remove traces of HFIP and stored dry at *−*20 ^*°*^C.

## ThT fluorescence assay

Aggregation kinetics were studied by tracking the rise in Thioflavin-T (ThT) fluorescence on a Victor X4 microplate reader (PerkinElmer). Measurements were performed in triplicate in a 96-well plate. ThT was excited at 450 nm and emission was detected at 490 nm. When indicated, normalized traces were analyzed using an AmyloFit-inspired numerical implementation of mechanistic aggregation models.^29^ The working model used for the DNAJB6-CTD experiments tracked the free monomer concentration, *m*(*t*), the aggregated monomer mass concentration, *M* (*t*), and the aggregate number concentration, *P* (*t*):

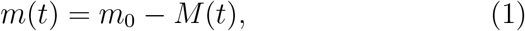

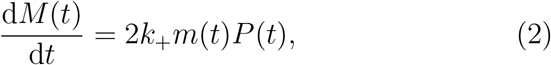

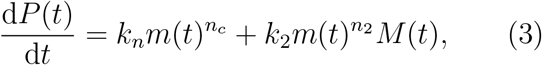

with *M* (0) = *P* (0) = 0. Here, *k*_*n*_, *k*_+_, and *k*_2_ are the apparent rate constants for primary nucleation, fibril-end elongation, and secondary nucleation, respectively. The reaction orders were fixed at *n*_*c*_ = *n*_2_ = 2. ThT fluorescence was assumed to be proportional to *M* (*t*), and the simulated aggregate-mass curves were normalized to their endpoint before comparison with the experimental traces. Following control-model comparison, the standard secondary-nucleation model was used as the working model for the DNAJB6-CTD analysis. The pooled control parameters were held fixed, and each DNAJB6-CTD condition was analyzed by allowing one rate parameter, *k*_*n*_, *k*_+_, or *k*_2_, to vary at a time.

### Cryo-electron microscopy

Samples were allowed to aggregate for 7 days at room temperature under quiescent conditions. Quantifoil 3.5/1 grids were glow-discharged for 90 s (20 mA), after which 3 *µ*L of sample was applied and vitrified by plunge-freezing into liquid ethane using a Vitrobot Mark IV (Thermo Scientific). Cryo-EM data were acquired at the Cryo-EM Uppsala facility on a 200 kV Glacios cryo-TEM equipped with a Falcon 4 direct electron detector. For each data set, 1000 movies were recorded using EPU Multigrid. Images were collected at a nominal magnification of 79,000*×* (pixel size 1.52 Å/px) with a target defocus range of *−*2.6 to *−*1.3 *µ*m. Each movie comprised 41 frames, corresponding to a total exposure of 57 e/Å^2^.

Unless otherwise stated, all processing steps were carried out identically for all data sets on the Tetralith and Arrhenius clusters at Sweden’s National Supercomputer Centre (NSC). Motion correction was performed in RELION 5.1^39–41^ using the internal implementation, followed by CTF estimation with CTFFIND-4.1. ^42^ Filaments were picked using the 3D-EM implementation of Topaz^43^ for fibrils: an autopicker model was trained on 100 manually annotated micrographs and subsequently used to automatically determine fibril start–end coordinates across the full data set. Particles were extracted using a 512-pixel box with an interbox distance corresponding to three asymmetric repeats (4.75 Å per repeat). For large data sets, particles were extracted from micrographs downsampled two-fold to reduce memory requirements. Two rounds of 2D classification were performed in RELION. An initial run used 200 classes with *T* = 2 (EM algorithm; CTFs ignored until the first peak). Classes containing fibrils were retained and subjected to a second 2D classification with *T* = 4. Polymorph distributions were assessed from the resulting class averages by visual comparison, and crossover distances were measured in ImageJ.^44^

### 2D IR spectroscopy

A Ti:sapphire regenerative amplifier (SpitFire, Spectra-Physics) operating at 3 kHz pumped a dual optical parametric amplifier (TOPAS-Twins, Light Conversion). In the high-energy arm, mid-IR pump pulses centered at 6100 nm were generated via non-collinear difference-frequency generation (NDFG) between the signal and idler beams. Probe and reference pulses were produced the same way in the low-energy arm. The pump beam was sent through a mid-IR pulse shaper (PhaseTech)^45–47^ to generate a pair of time-delayed pump pulses, while the probe delay was controlled with a mechanical delay stage. All beams were focused onto the sample, and the probe and reference beams were spectrally dispersed with a Horiba iHR-320 spectrograph and detected using a 2*×*64-element MCT array detector. 2D IR interferograms were acquired with a 12 fs time step over a total scan window of 2400 fs. The data were denoised using the active noise-reduction approach introduced by Feng and co-workers.^48,49^ Following aggregation, samples were lyophilized and resuspended in D_2_O to a final concentration of 1 *m*M for 2D IR spectroscopy.

### Amide I exciton simulations

Static one-exciton absorption spectra were calculated for idealized parallel *β*-sheets using a coupled-oscillator Hamiltonian. ^32,35,50^ Each strand contained six Amide I sites, and the number of strands and helical twist were varied. Local transition dipoles were assigned from the C=O and C–N geometry using the AIM/Torii–Tasumi parametrization.^51,52^ Sequential intrastrand couplings were fixed at +0.8 cm^*−*1^, while all remaining Amide I pairs were treated using the full transition-dipole coupling expression without a distance cutoff. Uniform site frequencies of 1650 cm^*−*1^ were used.

The resulting Hamiltonians were diagonalized exactly, and absorption spectra were generated from the exciton oscillator strengths using Lorentzian broadening with a full-width at half-maximum (FWHM) of 5 cm^*−*1^.

## RESULTS

### Initial hIAPP concentration shifts fibril polymorphism and produces non-monotonic aggregation kinetics

We previously showed that the polymorph distribution of hIAPP fibrils is strongly influenced by solution composition, including type of buffer and presence of peptide-based inhibitors. ^38^ Under low-ionic-strength conditions in 20 mM Tris buffer, 14 *µ*M hIAPP formed a nearly homogeneous population of short-crossover fibrils, which correspond to the previously reported double-S (TW3 polymorph),^24^ However, the effect of initial monomer concentration on the polymorph distribution under otherwise identical solution conditions had not been evaluated.

Here, we measured ThT aggregation kinetics at initial hIAPP concentrations of 14, 25, 50, 75, and 100 *µ*M in 20 mM Tris buffer (Figure 1A).^53^ Although all conditions produced sigmoidal fluorescence traces, the concentration dependence was non-monotonic. The time required to reach 50% of the maximum ThT signal, *t*_50_, was approximately 3.77, 4.62, 5.82, 5.41, and 5.15 h, respectively. Thus, *t*_50_ increased by 54% between 14 and 50 *µ*M before decreasing by 12% between 50 and 100 *µ*M. This behavior reveals two opposing concentration regimes with an apparent maximum near 50 *µ*M, rather than the expected monotonic acceleration at increasing monomer concentration. A similar critical-concentration dependence was reported by Brender et al., where they proposed a change in aggregation mechanism associated with micelle-like IAPP oligomers.^54^

**Figure 1.**
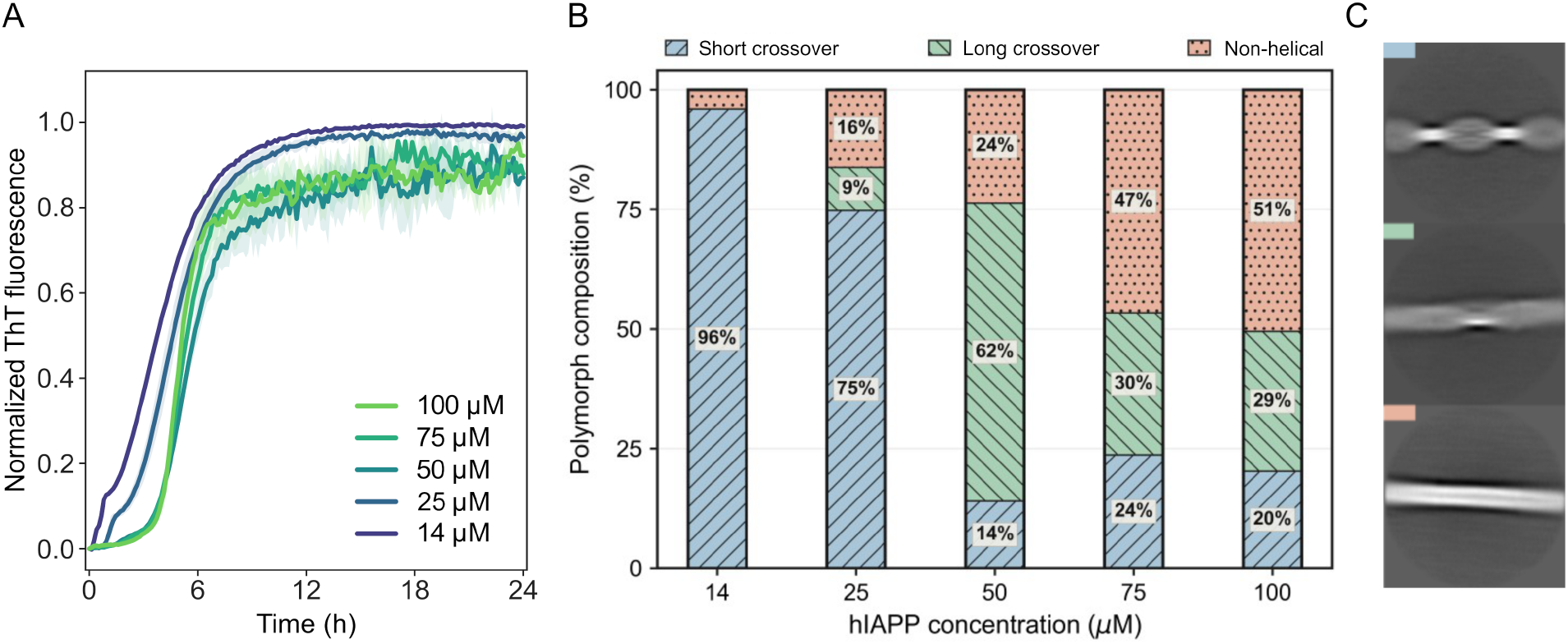
Concentration-dependent aggregation kinetics and fibril polymorphism of hIAPP in Tris buffer. (a) Normalized ThT fluorescence traces for hIAPP at concentrations of 14, 20, 50, 75, and 100 *µ*M in 20 mM Tris buffer. The traces are colored from dark blue for the lowest concentration to light green for the highest concentration. (b) Relative abundance of short crossover (pastel blue), long crossover (pastel green), and non-helical (pastel orange) fibrils in samples containing 14, 25, 50, 75, and 100 *µ*M hIAPP, determined by RELION 2D classification of cryo-EM particle segments. Data for the 14 *µ*M condition were taken from our previous study. ^38^ (c) Representative 2D class averages for each fibril morphology. Colored rectangles in the upper left corners identify the corresponding classes in panel (b).

The unexpected behavior in Tris was reproduced using two independently purified hIAPP batches (Figure S1A). In contrast, hIAPP aggregation in PBS showed the expected concentration dependence, with faster aggregation at higher peptide concentrations (Figure S1B). Thus, the unusual kinetic profile is specific to the low-ionic-strength Tris conditions. However, because ThT fluorescence reflects dye binding rather than secondary structure, these kinetic traces cannot distinguish changes in fibril structure. ^53^ We therefore proceeded to use cryo-EM to determine whether the endpoint fibril structures changed with initial hIAPP concentration.

Samples containing 25, 50, 75, and 100 *µ*M hIAPP were aggregated in Tris 20 mM buffer and analyzed by particle classification in RELION. ^41^ Fibrils were assigned to three morphological classes: short crossover fibrils, long crossover fibrils, and non-helical, flat fibrils of unknown structure. Representative class averages are presented in Figure 1C. Unlike the ThT kinetics, the cryo-EM analysis revealed a clear concentration-dependent redistribution of fibril morphologies (Figure 1B). At 14 *µ*M, short crossover fibrils accounted for 96% of the population, with only 4% non-helical fibrils and no detectable long crossover fibrils. Short crossover fibrils remained predominant at 25 *µ*M, but both long-crossover and non-helical fibrils appeared. At 50 *µ*M, long-crossover fibrils became the dominant class, accounting for 62% of the population. At 75 and 100 *µ*M, non-helical fibrils formed the largest class, increasing to 47% and 51%, respectively. Thus, increasing the initial hIAPP concentration shifted the endpoint population from a nearly monomorphic short crossover state toward a more heterogeneous ensemble enriched in non-helical species.

### Label-free 2D IR spectroscopy resolves concentration-dependent hIAPP polymorphism

Having established solution conditions that modulate hIAPP polymorph distributions, we decided to test if the polymorphs could be distingushied with label-free 2D IR spectra, rather than relying solely on cryo-EM. We measured label-free 2D IR spectra of hIAPP fibrils formed at 25, 50, and 100 *µ*M. 2D IR spectroscopy is highly sensitive to transition-dipole coupling and vibrational delocalization within amyloid *β*-sheets, but previous polymorph-resolved measurements have generally relied on isotopic labeling.^33–35^ The concentration-dependent samples therefore provided a direct test of whether fibril morphology is encoded in the unlabeled Amide I band.

The 2D IR spectra and their diagonal slices are shown in Figures 2A and 2B. At 25 *µ*M, the spectrum was dominated by a strongly blue-shifted band centered at approximately 1637 cm^*−*1^, with a weaker lower-frequency contribution near 1620 cm^*−*1^. At 50 *µ*M, the maximum shifted to approximately 1625 cm^*−*1^, although a distinct high-frequency shoulder remained visible near 1637 cm^*−*1^. At 100 *µ*M, this high-frequency contribution was no longer resolved, and the spectrum was dominated by a single band centered near 1622 cm^*−*1^.

**Figure 2.**
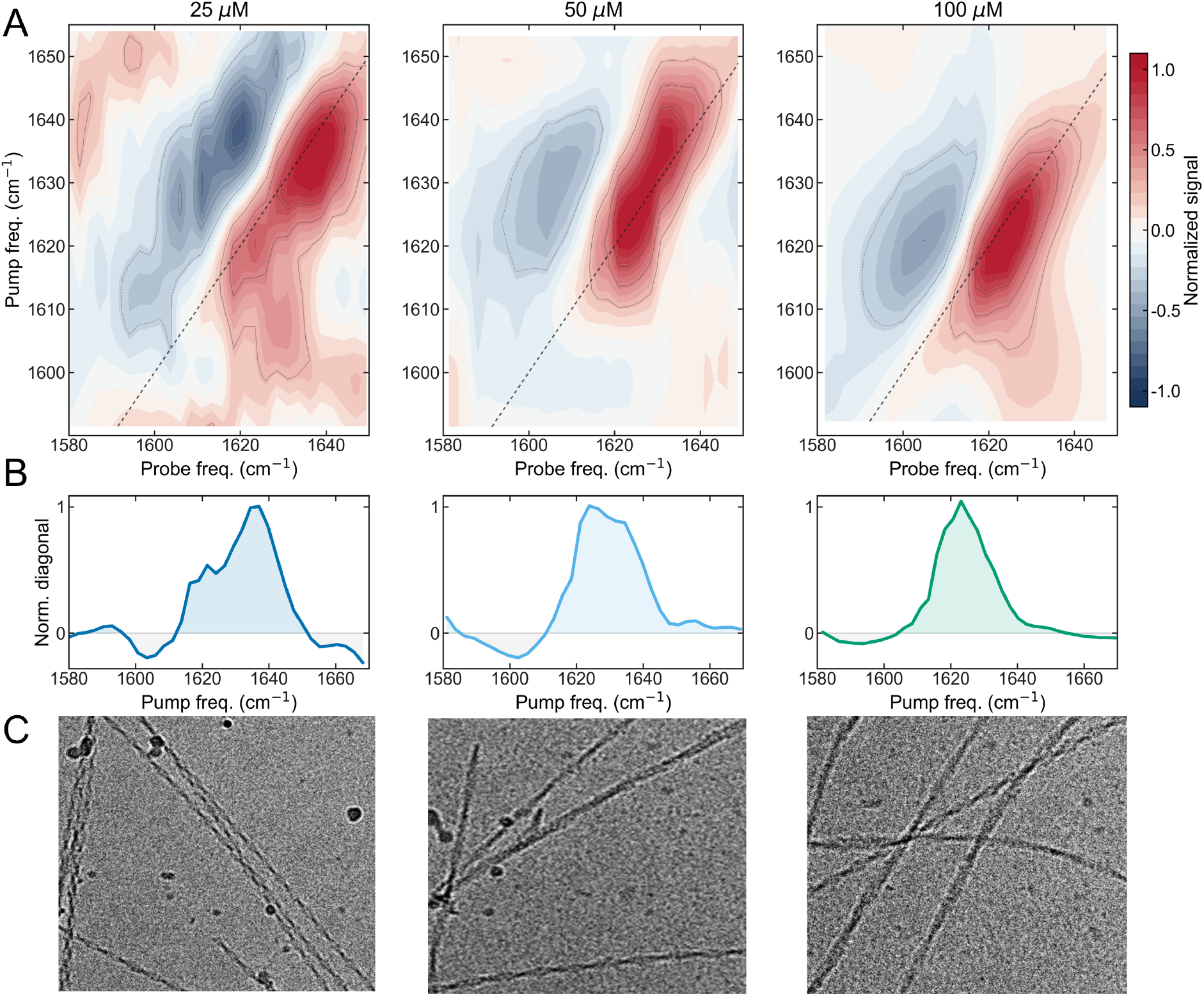
2D IR spectroscopy enables direct characterization of hIAPP amyloid polymorphism. (A) Representative 2D IR spectra of hIAPP aggregated in Tris buffer at peptide concentrations of 25, 50, and 100 *µ*M. (b) Diagonal slices extracted from the 2D IR spectra shown in (A), corresponding to hIAPP concentrations of 25, 50, and 100 *µ*M (left to right). (C) Representative cryo-EM micrographs illustrating the dependence of fibril morphology on initial peptide concentration: lower concentrations (25 *µ*M) are enriched in highly twisted fibrils, intermediate concentrations show a mixed population with emerging long crossover fibrils, and higher concentrations are dominated by long-crossover and non-helical fibrils.

This spectral progression closely followed the fibril distributions observed by cryo-EM (Figure 2C). The 25 *µ*M sample, which was enriched in short crossover fibrils, exhibited the pronounced 1637 cm^*−*1^ feature. The intermediate spectrum at 50 *µ*M accompanied the emergence of long crossover fibrils, whereas the low-frequency spectrum at 100 *µ*M showed the greatest abundance of non-helical fibrils. The high-frequency band is particularly interesting because it lies approximately 17 cm^*−*1^ above the Amide I frequency commonly associated with amyloid *β*-sheets.^35^ These results identify the 1637 cm^*−*1^ feature as a distinctive marker of short crossover-rich hIAPP fibril populations and demonstrate that label-free 2D IR spectroscopy can distinguish concentration-dependent polymorph distributions.

### Spectral simulations identify exciton-branch redistribution associated with helical twist

The Amide I response of a protein is dominated by stretching of the peptide-bond carbonyl groups. Interactions between these local vibrations produce collective excitonic states that extend over several amide groups.^32^ The frequency of each exciton is determined by the local Amide I frequencies and the couplings between them, whereas the intensities depend on the relative orientations and phases of the contributing transition dipoles. The diagonal 2D IR response therefore reflects both the energies of the excitonic states and the distribution of intensity among them. We used spectral simulations to determine how these quantities are affected by the helical geometry of the TW3 fibril.

We calculated the Amide I spectra of idealized *β*-sheets containing an increasing number of *β*-strands (Figure 3A). The positions and relative intensities change progressively with the nubmer of *β*-strands and are close to saturation point around the length of 20-25 strands. The previously reported convergence at 12-15 strands were due to a cut-off of 0.4 cm^*−*1^ applied to couplings, which is not used in our study. ^35^ A series of helical twists was then applied to the 20 strands *β*-sheet model (Figure 3B). Increasing the twist produced a clear redistribution of intensity between the low- (1610-1618 cm^*−*1^) and high-frequency (*>* 1618 cm^*−*1^) excitons. The high-frequency features became progressively stronger, accompanied by a reduction in the relative intensity of the low-frequency region. In comparison, the frequency of the brightest low-energy exciton changed only modestly across the twist series. The main calculated effect of twisting the ideal sheet was therefore a change in the relative transition strengths of the excitonic states, rather than a large frequency shift of the strongest low-energy exciton.

**Figure 3.**
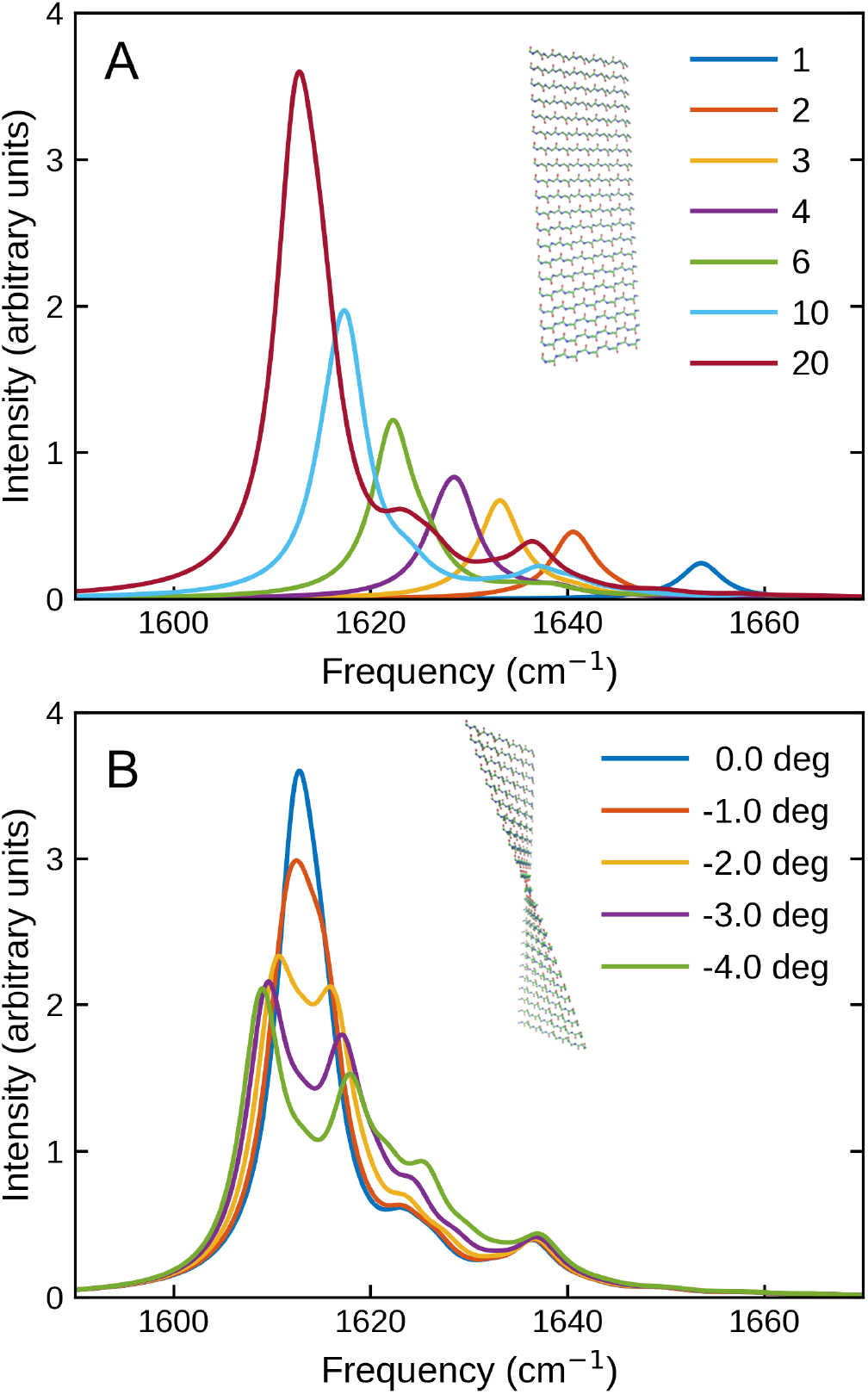
Spectral simulations of Amide I spectra identify exciton-branch redistribution associated with helical twist. (A) Calculated Amide I spectra for idealized beta-sheets containing an increasing number of beta-strands (from 1 - blue to 20 dark red), demonstrating convergence and near-saturation at approximately 20 to 25 strands. (B) Calculated spectra for a 20-strand beta-sheet model subjected to varying degrees of helical twist. Increasing the helical twist produces a clear redistribution of spectral intensity towards the higher frequencies

### Surface-associated nucleation can generate a fibril morphology distinct from its parent

Visual inspection of the cryo-EM micrographs of hIAPP revealed several branch-like fibril arrangements consistent with nucleation occurring on the surface of a pre-existing fibril (Figure 4). In the clearest example, a daughter fibril branches directly from a parent fibril displaying the short-crossover, double-S fold (Figure 4B). Notably, while the parent fibril exhibits a distinct periodic twist, the daughter fibril lacks detectable helical pitch and appears completely flat. Although a static cryo-EM image cannot establish the direction of growth or completely exclude an incidental fibril crossing, the branch-like geometry is consistent with a surface-catalyzed nucleation event.

**Figure 4.**
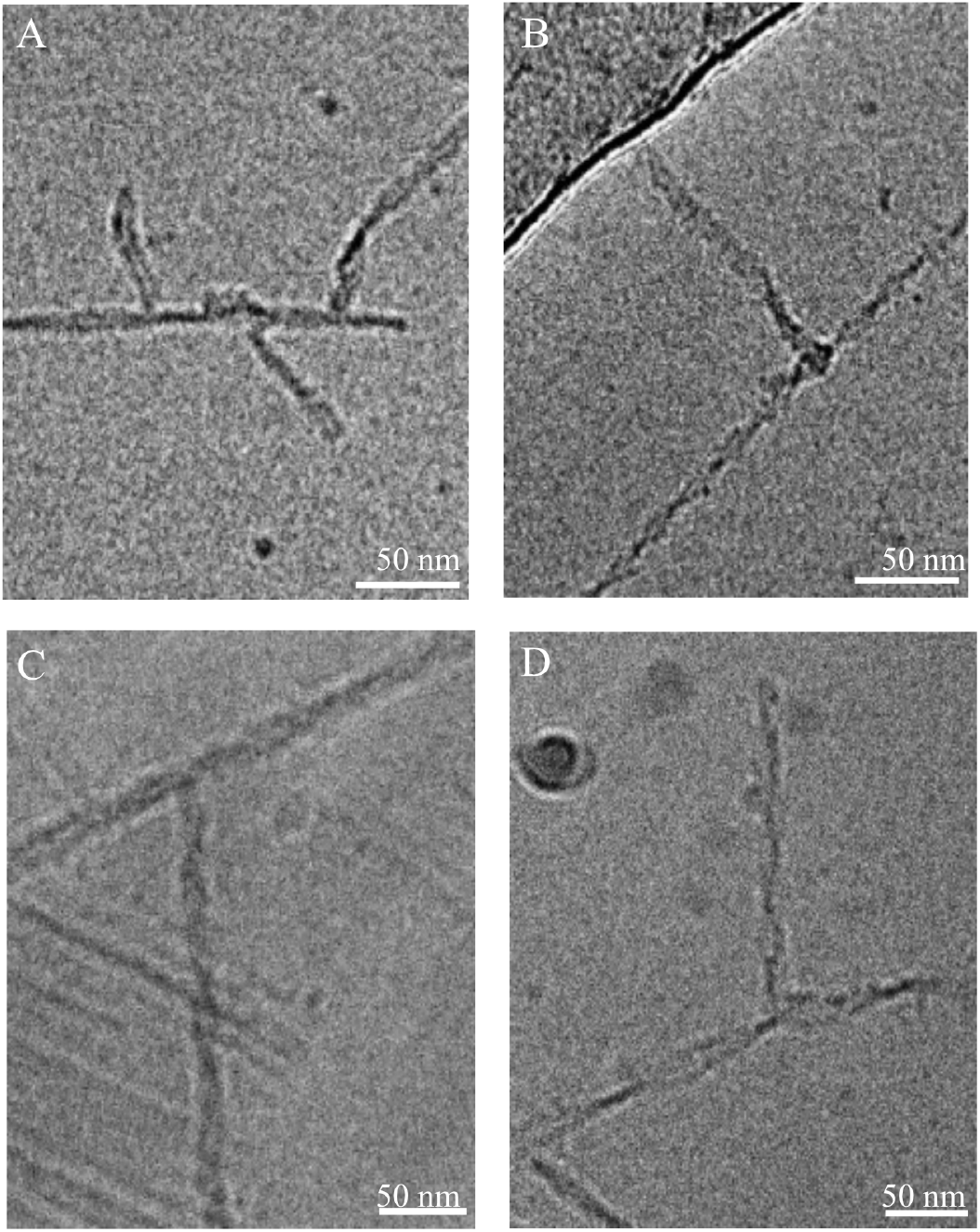
Cryo-EM images of secondary-nucleation events in hIAPP samples. (A) Multiple fibrils are associated with the surface of a pre-existing fibril. (B) A fibril emerges from a short crossover parent fibril consistent with the double-S polymorph. The parent exhibits a periodic crossover, whereas the associated fibril appears non-helical. (C, D) The reverse phenomenon can also occur, where a parent fibril exhibiting a long crossover nucleates the formation of a fibril with a shorter crossover on its surface.

This observation suggests that secondary nucleation may not always replicate the structure of the fibril that catalyzes it. Instead, the parent surface may promote the formation of a structurally distinct nucleus, providing a plausible mechanism for the increased polymorphic diversity observed at high hIAPP concentrations. This result motivated us to test whether selective suppression of secondary nucleation could redirect the final fibril distribution toward a more homogeneous state.

### DNAJB6-CTD suppresses secondary nucleation and shifts hIAPP fibril polymorphism

To test whether secondary nucleation drives this structural diversification, we perturbed this pathway using the isolated DNAJB6-CTD. This predominantly *β*-sheet domain has previously been shown to bind amyloid fibril surfaces and act as a selective inhibitor of secondary nucleation (Figure 5A).^36^

**Figure 5.**
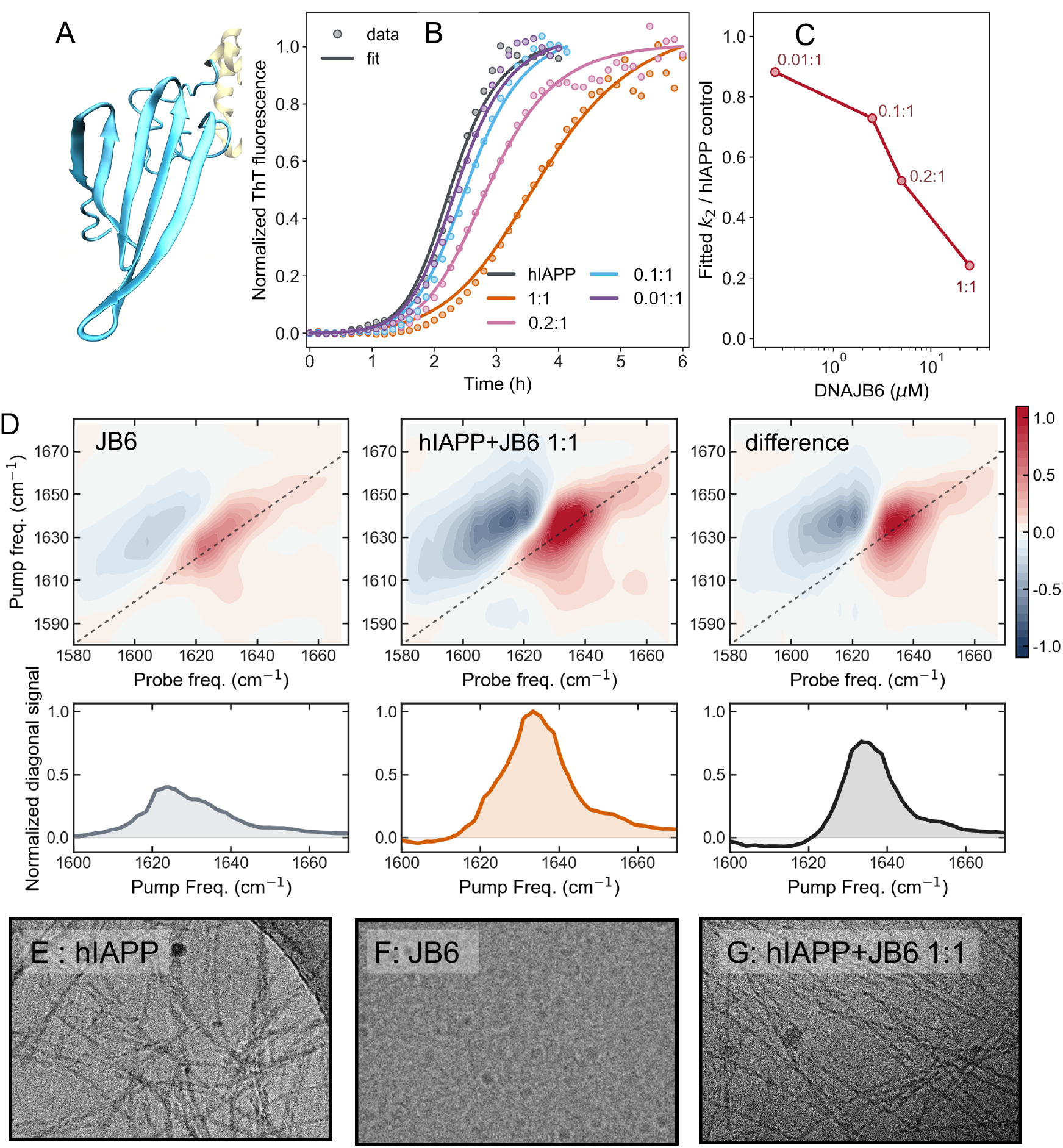
DNAJB6-CTD inhibits hIAPP secondary nucleation and alters the resulting fibril morphology. (A) Structure of the *β*-sheet-rich DNAJB6-CTD. The structure is prepared based on PDB ID:6U3R ^55^ using VMD softwere. ^56^ (B) Normalized ThT aggregation kinetics of 25 *µ*M hIAPP in PBS at the indicated CTD:hIAPP molar ratios. (C) Fitted secondary-nucleation rate constant, *k*_2_, normalized to the hIAPP-only control. (D) From left to right: 2D IR spectrum and diagonal slice of DNAJB6-CTD alone, the 1:1 CTD:hIAPP sample, and the difference spectrum obtained by subtracting the CTD-only spectrum from the mixed sample. (E-G) Representative cryo-EM micrographs of hIAPP alone, DNAJB6-CTD alone, and the 1:1 CTD:hIAPP sample, respectively.

Aggregation of 25 *µ*M hIAPP in PBS was monitored in the presence of DNAJB6-CTD at CTD:hIAPP molar ratios ranging from 0.01:1 to 1:1 (Figure 5B). Increasing DNAJB6-CTD concentration progressively delayed aggregation, with *t*_1*/*2_ increasing from 2.4 h for hIAPP alone to 3.8 h at a 1:1 ratio. To assess which microscopic process was most affected, the data were fit with an AmyloFit-inspired kinetic model^29^ containing primary nucleation, elongation, and secondary nucleation where only one rate constant was varied between datasets for different DNAJB6:hIAPP ratios (Figures 5C and S2). In this analysis, inhibition was captured primarily by a dose-dependent decrease in the fitted secondary-nucleation parameter, *k*_2_ (Figure 5C). Relative to hIAPP alone, the apparent *k*_2_ decreased from approximately 0.88 at a 0.01:1 ratio to 0.24 at a 1:1 ratio. These results are consistent with concentration-dependent suppression of hIAPP secondary-nucleation activity by DNAJB6-CTD.

To determine whether suppression of secondary nucleation altered the resulting hIAPP structures, we analyzed the 1:1 CTD:hIAPP sample by 2D IR spectroscopy (Figure 5D). Because DNAJB6-CTD contributes its own Amide I response, we measured the CTD-only spectrum and subtracted it from the spectrum of the mixed sample. The resulting difference spectrum exhibited a maximum near 1637 cm^*−*1^, matching the high-frequency feature observed for the short crossover-rich hIAPP sample in Figure 2.

Cryo-EM provided an independent test of this interpretation (Figures 5E–G). The hIAPP-only sample consisted of by non-helical fibrils, whereas the 1:1 CTD:hIAPP sample was dominated by short-crossover fibrils. DNAJB6-CTD alone formed small globular particles rather than fibrils. The agreement between the 1637 cm^*−*1^ difference signal and the short-crossover morphology therefore supports a shift toward the short crossover fibril class.

As an additional control, DNAJB6-CTD was incubated either with 10% hIAPP seeds or with 10% sonicated hIAPP seeds. Neither condition produced an increase in ThT fluorescence (Figure S3), indicating that hIAPP did not induce detectable ThT-positive CTD fibrillation under the conditions tested. Together, the kinetic and structural data support a model in which DNAJB6-CTD suppresses surface-catalyzed secondary nucleation and redirects hIAPP assembly toward short-crossover fibrils.

## DISCUSSION

### Concentration modulates hIAPP fibril polymorphism through kinetic partitioning

The hIAPP concentration series identify initial peptide concentration as a direct control parameter for hIAPP fibril polymorphism in Tris buffer. Under otherwise identical buffer conditions, the population of short-crossover fibrils changes from 96% at 14 *µ*M, via long crossover dominated ensemble at 50 *µ*M, towards primarily non-helical structures at 75-100 *µ*M. Thus, variation of a single experimental parameter was sufficient to generate a continuous redistribution of fibril morphologies without changing IAPP sequence, buffer, or incubation protocol.

The accompanying ThT kinetics do not show the monotonic acceleration normally expected when increasing monomer concentration. We identify two concentration regimes consistent with the previously reported NMR study which attribute this to the formation of micelle-like assemblies.^54^ Here, we present that this behaviour is specific to low ionic strength buffer media, as introducing PBS restores the trend in which higher hIAPP concentration results in faster aggregation. The present behavior is thus consistent with the strong sensitivity of hIAPP aggregation to buffer, ion identity, and ionic strength reported before. ^38,57–59^ In particular, a study of the hIAPP(11–20) segment found stronger association with Tris than with the other tested buffers, resulting in slower self-aggregation, whereas phosphate accelerated aggregation through electrostatic screening. ^57^

The most plausible explanation is that Tris shifts the concentration-dependent partitioning between productive and slowly converting peptide assemblies. Such behavior is expected when a competing off-pathway state becomes increasingly populated at high protein concentration. Kinetic modeling predicts that trapping monomers in these assemblies can reverse the usual concentration dependence of fibrillation, and this effect has been observed experimentally for ribosomal protein S6.^60,61^

At higher concentrations of hIAPP, the increasing rates of productive nucleation, elongation, or secondary processes could partially overcome this monomer depletion and account for the observed reversal above 50 *µ*M. However, the current ThT measurements do not distinguish among specific oligomerization and conversion steps. The DNAJB6-CTD experiments were thus used to gain further mechanistic insights into the role of secondary nucleation in polymorph formation.

### Imperfect templating driven by secondary nucleation shapes hIAPP polymorph distribution

DNAJB6-CTD has previously been shown to bind amyloid fibrils and suppress secondary nucleation of A*β*42.^36^ The present results extend this activity to hIAPP and establish DNAJB6-CTD as a useful perturbation for connecting aggregation kinetics with structural outcome. Models containing an autocatalytic fibril-multiplication pathway described the uninhibited hIAPP kinetics substantially better than a model containing only primary nucleation and elongation. Within the selected autocatalytic model, the DNAJB6-dependent delay was represented by a progressive decrease in the apparent secondary-nucleation parameter, *k*_2_. Because the normalized kinetic traces were collected at a single hIAPP concentration, a simultaneous effect on elongation cannot be excluded. Even with this limitation, the kinetic response, the established fibril-surface activity of DNAJB6-CTD, and the accompanying shift in fibril morphology support the interpretation that DNAJB6-CTD suppresses secondary nucleation during hIAPP aggregation.

The structural consequence of the inhibition was evident in the 2D IR spectra. After subtraction of the DNAJB6-CTD contribution, the mixed sample displayed the 1637 cm^*−*1^ feature associated with short-crossover TW3 polymorph. Cryo-EM independently confirmed enrichment of short-crossover fibrils relative to the heterogeneous hIAPP control. These results show that a molecular chaperone can influence amyloid structure by changing the relative contribution of competing microscopic pathways and 2D IR can provide a rapid detection of the resulting changes in fibril polymorphism.

The preferential enrichment of the TW3 short crossover polymorph is particularly interesting. TW3 contains folds related to those in the recently determined Ω-shaped fold isolated directly from pancreatic tissue of individuals with type 2 diabetes.^24,25^ Although TW3 and the ex vivo Ω fold are not identical, the recurrence of related motifs across multiple hIAPP structures suggests that this region of conformational space is especially accessible or stable. Suppression of secondary-pathway activity may therefore expose a structural basin that is efficiently reached during primary nucleation and faithfully propagated by fibril-end elongation, while secondary nucleation opens additional routes toward alternative architectures.

The results distinguish templating at fibril ends from nucleation on fibril surfaces. During elongation, the parent fibril end constrains H-bond registry, side-chain packing, and protofilament geometry, which favours faithful propagation of the existing architecture.^9,11^ A catalytic site on the lateral fibril surface provides a different set of constraints. Surface association can increase the local peptide concentration, restrict orientational freedom, and lower the barrier to forming an ordered nucleus; yet it need not reproduce the complete interaction network available at the fibril end. We refer to this process as imperfect templating. The parent fibril constrains the initial peptide arrangement through a partially structured interface, while the remaining contacts are selected during formation of the daughter nucleus. The daughter fibril can consequently retain some structural features favored by the parent surface while adopting a different overall morphology.

Recent evidence that secondary nucleation is concentrated at rare fibril growth defects provides a concrete molecular description for this model. Such defects can expose only part of the normally buried cross-*β* core, producing a highly active nucleation site without presenting the complete fibril-end interface.^62^ A partially exposed cross-section is precisely the type of scaffold expected to promote nucleation while allowing structural divergence. This defect-based picture also explains why catalytic activity may not always be distributed uniformly over the entire fibril surface.

The proposed mechanism aligns with evidence that surface templating can range from close replication of the parent structure to complete diversification, depending on the amyloid system. ^63^ For hIAPP, previous studies provide independent kinetic evidence for this diversifying mechanism: using isotopically labeled hIAPP, Farrell et al. tracked individual 2D IR cross-peaks and found that distinct polymorphs exhibited unique aggregation kinetics that could only be modeled by including secondary nucleation.^33^ This supports the formation of a daughter structure that structurally diverges from the parent template. Furthermore, direct imaging of A*β*-42 has captured daughter aggregates forming and detaching from parent fibril surfaces, confirming the physical geometry of surface-associated nucleation.^63^ The branch-like features we observe in our hIAPP cryo-EM micrographs are remarkably consistent with this geometry. In our clearest example, a non-helical fibril emerges directly from the lateral surface of a short crossover parent fibril (Figure 4). While these static images cannot definitively establish the timing or direction of growth, they are in agreement with our DNAJB6-CTD kinetic analyses. A secondary-nucleation pathway that occasionally produces a structurally distinct daughter fibril can make that morphology abundant, even if it is not the most thermodynamically stable structure, because each daughter nucleus generates a new fibril that can subsequently elongate and contribute additional catalytic sites.

### Polymorph-sensitive signatures in label-free 2D IR spectra

The ability of 2D IR spectroscopy to distinguish polymorphs from absorptive spectra alone without introducing isotope labels is unprecedented. Interestingly, the center frequency of the diagonal slice is a sufficient metric in our study to confidently assign the fibril structures to short-crossover vs non-helical fibrils. We are currently exploring possibilities for augmenting such analyses with transition dipole spectra, polarization control, and cross-peaks. Moreover, the possibility of forming a monomorphic distribution of fibrils with the aid of DNAJB6 allows for the collection of reference spectra specific to particular polymorphs found on cryo-EM grids. Such a library of polymorphs will enable rapid screening of fibril samples prior to costly and labor-intensive microscopy screening. This includes the ability both to quantify the relative abundance of specific polymorphs and to detect new, previously unseen structures.

Nevertheless, it remains to be determined why the 2D IR spectra of the twisted polymorph are so distinct from those of other, less-helical architectures. Based on our measurements of many different amyloid proteins, helical fibrils are consistently associated with much higher center frequencies, with some appearing as high as 1640 cm^*−*1^. Such high-frequency bands can be misinterpreted as non-amyloid signals because amyloid fibril bands are generally expected to appear in the range of 1610–1625 cm^*−*1^. Our simulations confirm that introducing a twist does weaken the adjacent-layer stacking couplings and transfers oscillator strength toward higher-frequency excitonic branches. However, the calculated shift of the spectral centroid is negligible, indicating that the average backbone twist angle alone does not fully account for the experimental shift in frequency. To account for a shift of this magnitude purely through changes in geometry, the distances between layers would have to be unrealistically large.

Thus, the spectral difference is unlikely to arise from twist alone. Other features associated with the twisted fold, such as changes in local packing, layer registry, hydration, electrostatics, or vibrational coupling to other modes, may also play a role. We foresee that improved theoretical models combined with cross-peak analysis will help develop more robust approaches to detecting specific fibril polymorphs.

## CONCLUSION

This study demonstrates that the targeted modulation of amyloid polymorphism serves as a powerful approach for elucidating the molecular mechanisms governing fibril assembly. The concentration-dependent redistribution from short crossover to non-helical fibrils shows that even a single experimental parameter can drive substantial structural evolution. Furthermore, the branch-like cryo-EM features provide direct visual evidence that secondary nucleation can generate daughter fibrils whose architectures diverge from their parent templates. This observation of imperfect templating on fibril surfaces challenges the view that amyloid propagation invariably preserves structural information and instead points to a diversification mechanism in which catalytic surfaces lower the barrier for nucleation without enforcing faithful replication.

Continued investigation into the parameters regulating polymorph selection may ultimately enable precise control over the structural ensembles generated *in vitro*. However, a significant challenge remains the narrow scope of environmental conditions under which this control is currently achievable. For instance, the concentration-dependent shift in polymorph distribution was uniquely observed in Tris buffer, whereas the selective inhibitory effect of the CTD domain was most pronounced in PBS. A critical outstanding question is whether pure populations of specific amyloid polymorphs can be selectively nucleated, rather than merely shifting the relative abundances within a heterogeneous mixture. Achieving absolute structural control will likely require the identification of specific solution conditions, sequence mutations, or molecular chaperones capable of reshaping the conformational energy landscape to strongly favor a single, thermodynamically preferred architecture. Such advances would represent a major step toward the deterministic control of amyloid aggregation.

The shift toward the TW3-related short crossover architecture upon chaperone treatment is particularly noteworthy, as this fold shares key structural motifs with the *ex vivo* hIAPP fibrils recently isolated from human pancreatic tissue. This convergence raises the intriguing possibility that suppression of secondary nucleation—whether by chaperones or by other environmental factors—may be a key determinant of the restricted polymorph landscapes observed in disease.

From a methodological standpoint, we managed to spectroscopically distinguish short crossover from flat polymorphs of hIAPP without isotopic labeling. This helps establishing 2D IR spectroscopy as a valuable complement to cryo-EM for high-throughput structural screening. The 1637 cm^*−*1^ feature we identify as a marker of twisted fibril populations provides a rapid detection of twisted polymorph that can guide the selection of conditions for subsequent microscopy. However, the physical origin of this unusually high-frequency Amide I band demands further analysis.

Taken together, our results identify secondary nucleation as both a source of amyloid structural diversification and an experimentally accessible control point for polymorph selection, providing a mechanistic basis for understanding—and ultimately directing—the structural outcomes of amyloid assembly.

## Acknowledgement

MM acknowledges the Swedish Society for Medical Research (S20-0156), Diabetesfonden (DIA2025-1003), and Magnus Bergvalls Foundation (grant no. 2025-415). The computations were enabled by resources provided by the National Academic Infrastructure for Supercomputing in Sweden (NAISS) at Uppsala University and Linköping University partially funded by the Swedish Research Council through grant agreement no. 2022-06725 (NAISS). Computing time was provided with the following grants: NAISS 2025/5-307, NAISS 2026/3-467, and NAISS 2026/3-89. For sample preparation, grid screening and data collection, we acknowledge the use of the Cryo-EM Uppsala facility, funded by the Department of Cell and Molecular Biology, and by the Disciplinary Domains of Science and Technology and of Medicine and Pharmacy at Uppsala University.

## Author contributions statement

MIK: Methodology, Investigation, Data curation, Visualization, Writing – original draft, Writing - review & editing, Conceptualisation. L-K: Investigation, Data curation, Visualization. SP Investigation, Data curation. DV: Investigation, Data curation. MM: Writing – review & editing, Writing – original draft, Visualization, Validation, Supervision, Software, Resources, Project administration, Methodology, Investigation, Funding acquisition, Formal analysis, Data curation, Conceptualization.

## Data availability statement

The data that support the findings of this study are available from the corresponding author upon reasonable request.

## Supplementary Information to: Secondary nucleation drives polymorph diversity in hIAPP amyloids

**Supplementary Figure 1:**
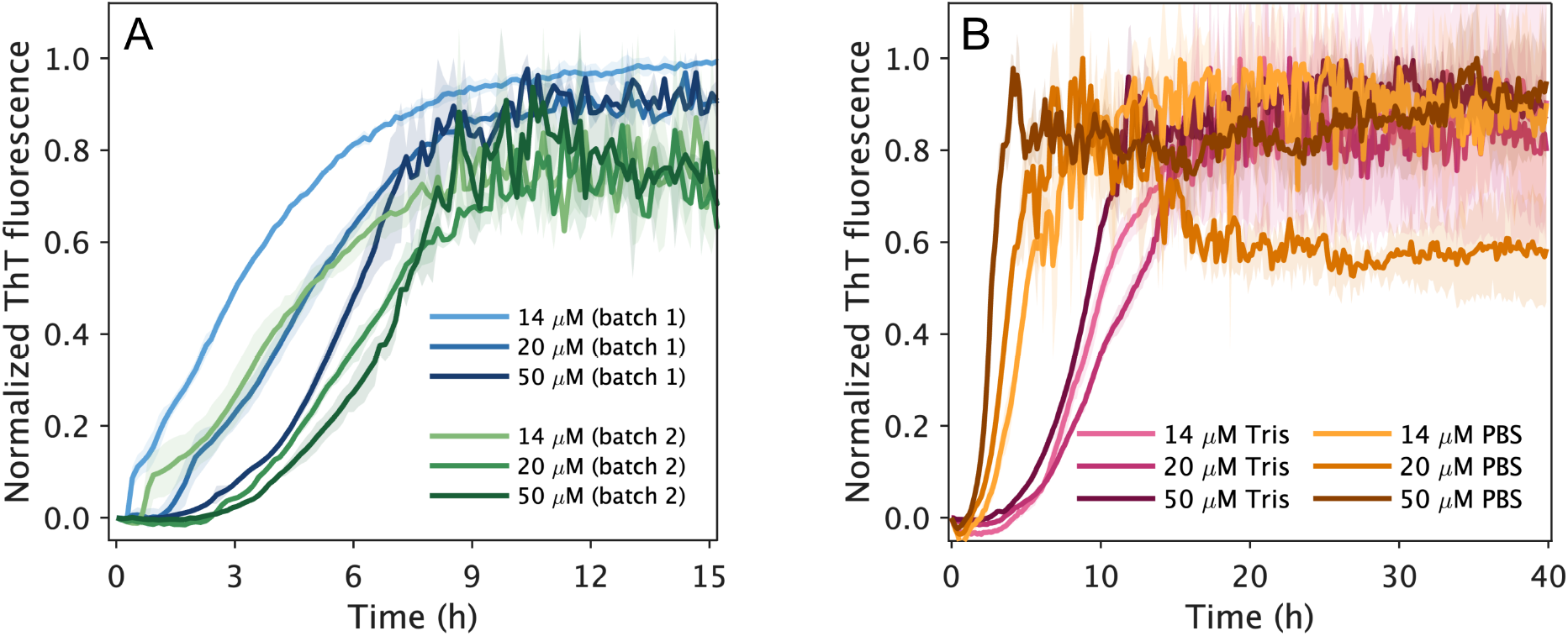
Aggregation kinetics extending the concentration-dependence experiments. A: Normalized ThT fluorescence traces of hIAPP at concentrations of 14, 20, and 50 µM in 20 mM Tris buffer. To assess the reproducibility of the non-canonical ThT kinetics observed in Tris buffer, experiments were repeated using independent peptide batches. The color gradient indicates increasing peptide concentration. B: Normalized ThT fluorescence traces illustrating the concentration dependence of aggregation kinetics in two different buffers. The pink color gradient represents three consecutive hIAPP concentrations (14, 20, and 50 µM) in Tris buffer, while the orange color gradient represents the corresponding concentrations in PBS buffer.

**Supplementary Figure 2:**
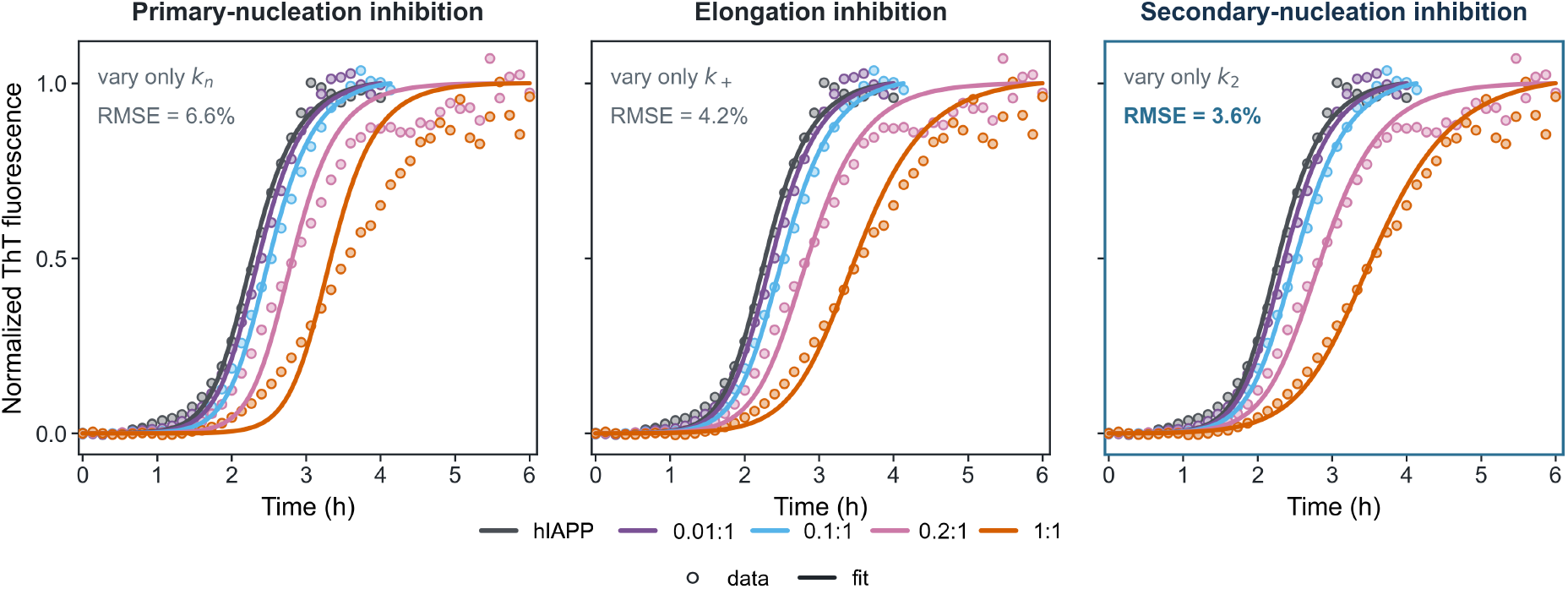
Secondary nucleation is inhibited by DNAJB6 in hIAPP aggregation. Experimental aggregation kinetics (points) were fitted using kinetic models in which only one rate constant was varied between datasets: the primary-nucleation rate constant, *k*_2_ (left); *k*_+_ the elongation rate constant, *k*_+_ (middle); or the secondary-nucleation rate constant, *k*_2_ (right). The corresponding root-mean-square errors (RMSEs) indicate that varying (*k*_2_) provides the best overall agreement with the experimental data (RMSE = 3.6%), compared with varying (*k*_+_) (4.2%) or (*k*_*n*_) (6.6%).

**Supplementary Figure 3:**
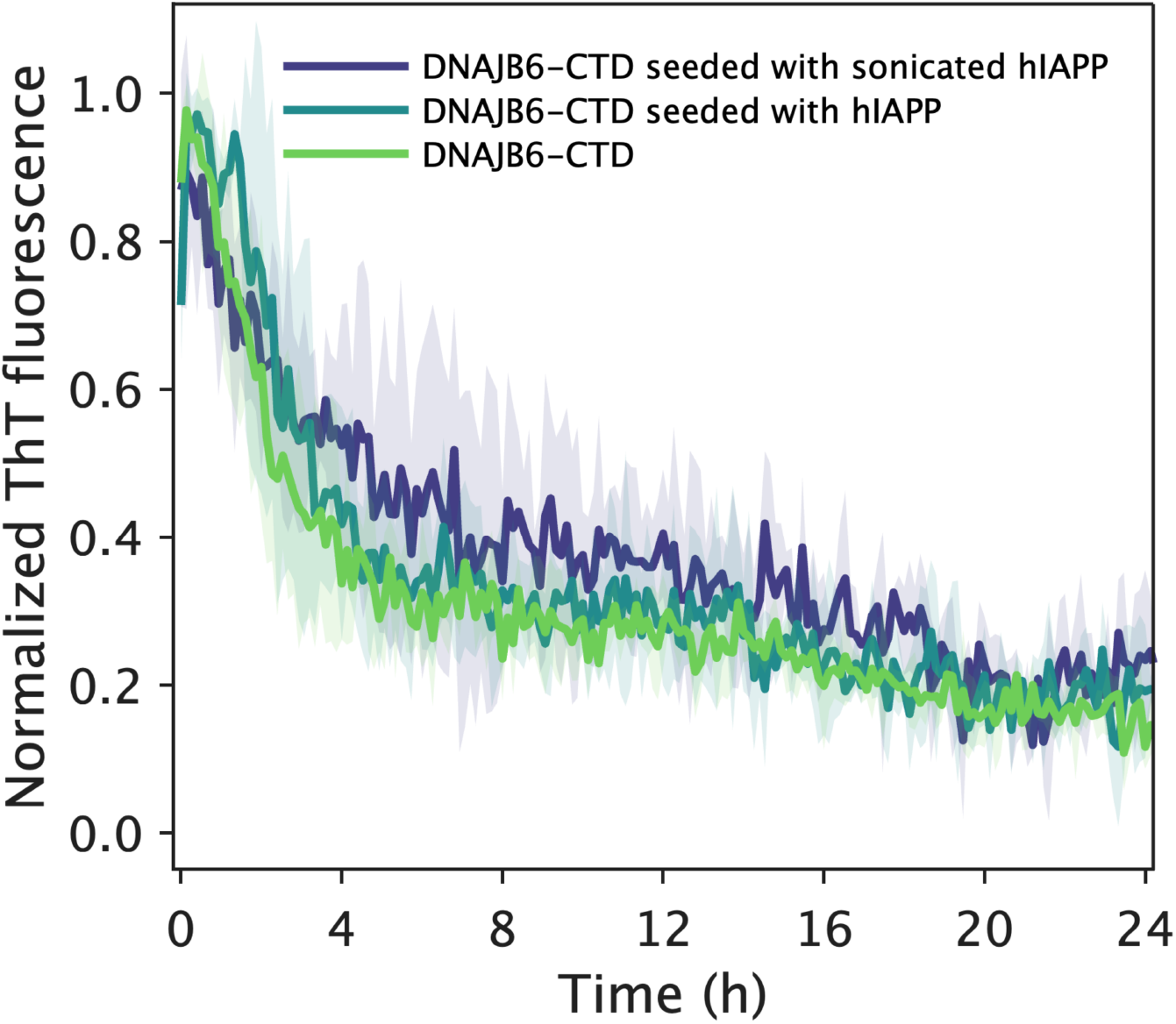
ThT aggregation kinetics of the DNAJB6 C-terminal domain (CTD) seeded with hIAPP fibrils. CTD was incubated at 25 µM in 20 mM Tris buffer either alone (light green) or in the presence of 10% (v/v) hIAPP seeds. Seeds were prepared either without sonication (teal) or after sonication (dark blue).

## References

(1) Chiti, F.; Dobson, C. M. Protein Misfolding, Functional Amyloid, and Human Disease. Annu. Rev. Biochem. 2006, 75, 333–366.

(2) Knowles, T. P. J.; Vendruscolo, M.; Dobson, C. M. The Amyloid State and Its Association with Protein Misfolding Diseases. Nat. Rev. Mol. Cell Biol. 2014, 15, 384–396.

(3) Hardy, J.; Selkoe, D. J. The Amyloid Hypothesis of Alzheimer’s Disease: Progress and Problems on the Road to Therapeutics. Science 2002, 297, 353–356.

(4) Haataja, L.; Gurlo, T.; Huang, C. J.; Butler, P. C. Islet Amyloid in Type 2 Diabetes, and the Toxic Oligomer Hypothesis. Endocr. Rev. 2008, 29, 303–316.

(5) Westermark, P.; Wernstedt, C.; Wilander, E.; Hayden, D. W.; O’Brien, T. D.; Johnson, K. H. Amyloid Fibrils in Human Insulinoma and Islets of Langerhans of the Diabetic Cat Are Derived from a Neuropeptide-Like Protein Also Present in Normal Islet Cells. Proc. Natl. Acad. Sci. U.S.A. 1987, 84, 3881–3885.

(6) Eisenberg, D.; Jucker, M. The Amyloid State of Proteins in Human Diseases. Cell 2012, 148, 1188–1203.

(7) Tycko, R. Amyloid Polymorphism: Structural Basis and Neurobiological Relevance. Annu. Rev. Biophys. 2015, 44, 1–23.

(8) Paravastu, A. K.; Leapman, R. D.; Yau, W.-M.; Tycko, R. Molecular Structural Basis for Polymorphism in Alzheimer’s β-Amyloid Fibrils. Proc. Natl. Acad. Sci. U.S.A. 2008, 105, 18349–18354.

(9) Petkova, A. T.; Leapman, R. D.; Guo, Z.; Yau, W.-M.; Mattson, M. P.; Tycko, R. Self-Propagating, Molecular-Level Polymorphism in Alzheimer’s β-Amyloid Fibrils. Science 2005, 307, 262–265.

(10) Bousset, L.; Pieri, L.; Ruiz-Arlandis, G.; Gath, J.; Jensen, P. H.; Habenstein, B.; Madiona, K.; Olieric, V.; Böckmann, A.; Meier, B. H.; Melki, R. Structural and Functional Characterization of Two α-Synuclein Strains. Nat. Commun. 2013, 4, 2575.

(11) Sanders, D. W.; Kaufman, S. K.; DeVos, S. L.; Sharma, A. M.; Mirbaha, H.; Li, A.; Barker, S. J.; Foley, A. C.; Thorpe, J. R.; Serpell, L. C.; Miller, T. M.; Grinberg, L. T.; Seeley, W. W.; Diamond, M. I. Distinct Tau Prion Strains Propagate in Cells and Mice and Define Different Tauopathies. Neuron 2014, 82, 1271–1288.

(12) Fitzpatrick, A. W. P.; Falcon, B.; He, S.; Murzin, A. G.; Murshudov, G.; Garringer, H. J.; Crowther, R. A.; Ghetti, B.; Goedert, M.; Scheres, S. H. W. Cryo-EM Structures of Tau Filaments from Alzheimer’s Disease. Nature 2017, 547, 185–190.

(13) Schweighauser, M.; Shi, Y.; Tarutani, A.; Kametani, F.; Murzin, A. G.; Ghetti, B.; Matsubara, T.; Tomita, T.; Ando, T.; Hasegawa, K.; Murayama, S.; Yoshida, M.; Hasegawa, M.; Scheres, S. H. W.; Goedert, M. Structures of α-Synuclein Filaments from Multiple System Atrophy. Nature 2020, 585, 464–469.

(14) Yang, Y. et al. Cryo-EM Structures of Amyloid-β 42 Filaments from Human Brains. Science 2022, 375, 167–172.

(15) Lövestam, S.; Koh, F. A.; van Knippenberg, B.; Kotecha, A.; Murzin, A. G.; Goedert, M.; Scheres, S. H. W. Assembly of Recombinant Tau into Filaments Identical to Those of Alzheimer’s Disease and Chronic Traumatic Encephalopathy. eLife 2022, 1, e76494.

(16) Zhang, W.; Falcon, B.; Murzin, A. G.; Fan, J.; Crowther, R. A.; Goedert, M.; Scheres, S. H. W. Heparin-Induced Tau Filaments Are Polymorphic and Differ from Those in Alzheimer’s and Pick’s Diseases. eLife 2019, 8, e43584.

(17) Falcon, B.; Zhang, W.; Schweighauser, M.; Murzin, A. G.; Vidal, R.; Garringer, H. J.; Ghetti, B.; Scheres, S. H. W.; Goedert, M. Tau Filaments from Multiple Cases of Sporadic and Inherited Alzheimer’s Disease Adopt a Common Fold. Acta Neuropathol. 2018, 136, 699–708.

(18) Kollmer, M.; Close, W.; Funk, L.; Bsoul, A.; Schmidt, M.; Rasmussen, J.; Jucker, M.; Fändrich, M. Cryo-EM Structure and Polymorphism of Aβ Amyloid Fibrils Purified from Alzheimer’s Brain Tissue. Nat. Commun. 2019, 10, 4760.

(19) Lövestam, S.; Schweighauser, M.; Matsubara, T.; Murayama, S.; Tomita, T.; Ando, T.; Hasegawa, K.; Yoshida, M.; Tarutani, A.; Hasegawa, M.; Goedert, M.; Scheres, S. H. W. Seeded Assembly in vitro Does Not Replicate the Structures of α-Synuclein Filaments from Multiple System Atrophy. FEBS Open Bio 2021, 11, 999–1013.

(20) Cao, Q.; Boyer, D. R.; Sawaya, M. R.; Ge, P.; Eisenberg, D. S. Cryo-EM Structure and Inhibitor Design of Human IAPP (Amylin) Fibrils. Nat. Struct. Mol. Biol. 2020, 27, 653–659.

(21) Röder, C.; Kupreichyk, T.; Gremer, L.; Schenk, C.; Fändrich, M.; Willbold, D.; Schröder, G. F. Cryo-EM Structure of Islet Amyloid Polypeptide Fibrils Reveals Similarities with Amyloid-β Fibrils. Nat. Struct. Mol. Biol. 2020, 27, 660–667.

(22) Gallardo, R.; Iadanza, M. G.; Xu, Y.; Heath, G. R.; Foster, R.; Radford, S. E.; Ranson, N. A. Fibril Structures of Diabetes-Related Amylin Variants Reveal a Basis for Surface-Templated Assembly. Nat. Struct. Mol. Biol. 2020, 27, 1048–1056.

(23) Li, D.; Zhang, X.; Wang, Y.; Zhang, H.; Song, K.; Bao, K.; Zhu, P. A New Polymorphism of Human Amylin Fibrils with Similar Protofilaments and a Conserved Core. iScience 2022, 25, 105705.

(24) Cao, Q.; Boyer, D. R.; Sawaya, M. R.; Abskharon, R.; Saelices, L.; Nguyen, B. A.; Lu, J.; Murray, K. A.; Kandeel, F.; Eisenberg, D. S. Cryo-EM Structures of hIAPP Fibrils Seeded by Patient-Extracted Fibrils Reveal New Polymorphs and Conserved Fibril Cores. Nat. Struct. Mol. Biol. 2021, 28, 724–730.

(25) Liu, W.; Han, J.; Gong, W.; Zhang, F.; Cao, Q. Structure of Pancreatic hIAPP Fibrils Derived from Patients with Type 2 Diabetes. Cell 2026, Online ahead of print (published Jan 2, 2026).

(26) Knowles, T. P. J.; Waudby, C. A.; Devlin, G. L.; Cohen, S. I. A.; Aguzzi, A.; Vendruscolo, M.; Terentjev, E. M.; Welland, M. E.; Dobson, C. M. An Analytical Solution to the Kinetics of Breakable Filament Assembly. Science 2009, 326, 1533–1537.

(27) Cohen, S. I. A.; Linse, S.; Luheshi, L. M.; Hellstrand, E.; White, D. A.; Rajah, L.; Otzen, D. E.; Vendruscolo, M.; Dobson, C. M.; Knowles, T. P. J. Proliferation of Amyloid-β42 Aggregates Occurs through a Secondary Nucleation Mechanism. Proc. Natl. Acad. Sci. U.S.A. 2013, 110, 9758–9763.

(28) Meisl, G.; Yang, X.; Hellstrand, E.; Frohm, B.; Kirkegaard, J. B.; Cohen, S. I. A.; Dobson, C. M.; Linse, S.; Knowles, T. P. J. Differences in Nucleation Behavior Underlie the Contrasting Aggregation Kinetics of the Aβ40 and Aβ42 Peptides. Proc. Natl. Acad. Sci. U.S.A. 2014, 111, 9384–9389.

(29) Meisl, G.; Kirkegaard, J. B.; Arosio, P.; Michaels, T. C. T.; Vendruscolo, M.; Dobson, C. M.; Linse, S.; Knowles, T. P. J. Molecular Mechanisms of Protein Aggregation from Global Fitting of Kinetic Models. Nat. Protoc. 2016, 11, 252–272.

(30) Rodriguez Camargo, D. C.; Chia, S.; Menzies, J.; Mannini, B.; Meisl, G.; Lundqvist, M.; Pohl, C.; Bernfur, K.; Lattanzi, V.; Habchi, J.; Cohen, S. I. A.; Knowles, T. P. J.; Vendruscolo, M.; Linse, S. Surface-Catalyzed Secondary Nucleation Dominates the Generation of Toxic IAPP Aggregates. Front. Mol. Biosci. 2021, 8, 757425.

(31) Wilkinson, M.; Xu, Y.; Thacker, D.; Taylor, A. P.; Fisher, D. G.; Gallardo, R. U.; Radford, S. E.; Ranson, N. A. Structural Evolution of Fibril Polymorphs during Amyloid Assembly. Cell 2023, 186, 5798–5811.e26.

(32) Hamm, P.; Zanni, M. Concepts and Methods of 2D Infrared Spectroscopy; Cambridge University Press: Cambridge, 2011.

(33) Farrell, K. M.; Fields, C. R.; Dicke, S. S.; Zanni, M. T. Simultaneously Measured Kinetics of Two Amyloid Polymorphs Using Cross Peak Specific 2D IR Spectroscopy. J. Phys. Chem. Lett.s 2023, 14, 11750–11757.

(34) Weeks, W. B.; Buchanan, L. E. Label-Free Detection of β-Sheet Polymorphism. J. Phys. Chem. Lett.s 2022, 13, 9449–9454.

(35) Lomont, J. P.; Ostrander, J. S.; Ho, J.-J.; Petti, M. K.; Zanni, M. T. Not All β-Sheets Are the Same: Amyloid Infrared Spectra, Transition Dipole Strengths, and Couplings Investigated by 2D IR Spectroscopy. J. Phys. Chem. B 2017, 121, 8935–8945.

(36) Österlund, N.; Frankel, R.; Carlsson, A.; Thacker, D.; Karlsson, M.; Matus, V.; Gräslund, A.; Emanuelsson, C.; Linse, S. The C-Terminal Domain of the Antiamyloid Chaperone DNAJB6 Binds to Amyloid-β Peptide Fibrils and Inhibits Secondary Nucleation. J. Biol. Chem. 2023, 299, 105317.

(37) Abedini, A.; Raleigh, D. P. Incorporation of Pseudoproline Derivatives Allows the Facile Synthesis of Human IAPP, a Highly Amyloidogenic and Aggregation-Prone Polypeptide. Org. Lett. 2005, 7, 693–696.

(38) Valli, D.; Ooi, S. A.; Kaya, I.; Thomassen, A. B.; Chaudhary, H.; Weidner, T.; Andrën, P. E.; Maj, M. Cryo-Electron Microscopy Provides Mechanistic Insights into Solution-Dependent Polymorphism and Cross-Aggregation Phenomena of the Human and Rat Islet Amyloid Polypeptides. Biochemistry 2025, 64, 2583–2595.

(39) Scheres, S. H. W. RELION: Implementation of a Bayesian Approach to Cryo-EM Structure Determination. J. Struct. Biol. 2012, 180, 519–530.

(40) Zivanov, J.; Nakane, T.; Forsberg, B. O.; Kimanius, D.; Hagen, W. J. H.; Lindahl, E.; Scheres, S. H. W. New Tools for Automated High-Resolution Cryo-EM Structure Determination in RELION-3. eLife 2018, 7, e42166.

(41) Kimanius, D.; Dong, L.; Sharov, G.; Nakane, T.; Scheres, S. H. W. New Tools for Automated Cryo-EM Single-Particle Analysis in RELION-4.0. Biochem. J. 2021, 478, 4169–4185.

(42) Rohou, A.; Grigorieff, N. CTFFIND4: Fast and Accurate Defocus Estimation from Electron Micrographs. J. Struct. Biol. 2015, 192, 216–221.

(43) Bepler, T.; Morin, A.; Rapp, M.; Brasch, J.; Shapiro, L.; Noble, A. J.; Berger, B. Positive-unlabeled Convolutional Neural Networks for Particle Picking in Cryo-Electron Microscopy. Nat. Methods 2019, 16, 1153–1160.

(44) Schneider, C. A.; Rasband, W. S.; Eliceiri, K. W. NIH Image to ImageJ: 25 years of image analysis. Nat. Methods 2012, 9, 671–675.

(45) Shim, S.-H.; Strasfeld, D. B.; Fulmer, E. C.; Zanni, M. T. Femtosecond Pulse Shaping Directly in the Mid-IR Using Acousto-Optic Modulation. Opt. Lett. 2006, 31, 838–840.

(46) Shim, S.-H.; Strasfeld, D. B.; Zanni, M. T. Generation and Characterization of Phase and Amplitude Shaped Femtosecond Mid-IR Pulses. Opt. Express 2006, 14, 13120–13130.

(47) Shim, S.-H.; Strasfeld, D. B.; Ling, Y. L.; Zanni, M. T. Automated 2D IR Spectroscopy Using a Mid-IR Pulse Shaper and Application of This Technology to the Human Islet Amyloid Polypeptide. Proc. Natl. Acad. Sci. U.S.A. 2007, 104, 14197–14202.

(48) Feng, Y.; Vinogradov, I.; Ge, N.-H. General noise suppression scheme with reference detection in heterodyne nonlinear spectroscopy. Opt. Express 2017, 25, 26262–26279.

(49) Feng, Y.; Vinogradov, I.; Ge, N.-H. Optimized noise reduction scheme for heterodyne spectroscopy using array detectors. Opt. Express 2019, 27, 20323–20338.

(50) Karjalainen, E.-L.; Ravi, H. K.; Barth, A. Simulation of the Amide I Absorption of Stacked β-Sheets. J. Phys. Chem. B 2011, 115, 749–757.

(51) van Adrichem, K. E.; Jansen, T. L. C. AIM: A Mapping Program for Infrared Spectroscopy of Proteins. J. Chem. Theory Comput. 2022, 18, 3089–3098.

(52) Torii, H.; Tasumi, M. Model Calculations on the Amide-I Infrared Bands of Globular Proteins. J. Chem. Phys. 1992, 96, 3379–3387.

(53) LeVine, H. Thioflavine T Interaction with Synthetic Alzheimer’s Disease β-Amyloid Peptides: Detection of Amyloid Aggregation in Solution. Protein Sci. 1993, 2, 404–410.

(54) Brender, J. R.; Krishnamoorthy, J.; Sciacca, M. F. M.; Vivekanandan, S.; D’Urso, L.; Chen, J.; La Rosa, C.; Ramamoorthy, A. Probing the Sources of the Apparent Irreproducibility of Amyloid Formation: Drastic Changes in Kinetics and a Switch in Mechanism Due to Micelle-Like Oligomer Formation at Critical Concentrations of IAPP. J. Phys. Chem. B 2015, 119, 2886–2896.

(55) Karamanos, T. K.; Tugarinov, V.; Clore, G. M. Unraveling the structure and dynamics of the human DNAJB6b chaperone by NMR reveals insights into Hsp40-mediated proteostasis. Proc. Natl. Acad. Sci. U.S.A. 2019, 116, 21529–21538.

(56) Humphrey, W.; Dalke, A.; Schulten, K. VMD: Visual molecular dynamics. J. Mol. Graph. 1996, 14, 33–38.

(57) Mao, Y.; Yu, L.; Yang, R.; Ma, C.; bo Qu, L.; de B. Harrington, P. New insights into side effect of solvents on the aggregation of human islet amyloid polypeptide 11–20. Talanta 2016, 148, 380–386.

(58) Marek, P. J.; Patsalo, V.; Green, D. F.; Raleigh, D. P. Ionic Strength Effects on Amyloid Formation by Amylin Are a Complicated Interplay among Debye Screening, Ion Selectivity, and Hofmeister Effects. Biochemistry 2012, 51, 8478–8490.

(59) Young, L. M.; Cao, P.; Raleigh, D. P.; Ashcroft, A. E.; Radford, S. E. Ion Mobility Spectrometry–Mass Spectrometry Defines the Oligomeric Intermediates in Amylin Amyloid Formation and the Mode of Action of Inhibitors. J. Am. Chem. Soc. 2014, 136, 660–670.

(60) Powers, E. T.; Powers, D. L. Mechanisms of Protein Fibril Formation: Nucleated Polymerization with Competing Off-Pathway Aggregation. Biophysical Journal 2008, 94, 379–391.

(61) Deva, T.; Lorenzen, N.; Vad, B. S.; Petersen, S. V.; Thørgersen, I.; Enghild, J. J.; Kristensen, T.; Otzen, D. Off-Pathway Aggregation Can Inhibit Fibrillation at High Protein Concentrations. Biochim. Biophys. Acta, Proteins Proteomics 2013, 1834, 677–687.

(62) Hu, J. et al. Structural Defects in Amyloid-β Fibrils Drive Secondary Nucleation. Nat. Commun. 2026, 17, 1933.

(63) Thacker, D.; Barghouth, M.; Bless, M.; Zhang, E.; Linse, S. Direct Observation of Secondary Nucleation along the Fibril Surface of the Amyloid-β 42 Peptide. Proc. Natl. Acad. Sci. U.S.A. 2023, 120,e2220664120.

